# EGAD: Ultra-fast functional analysis of gene networks

**DOI:** 10.1101/053868

**Authors:** Sara Ballouz, Melanie Weber, Paul Pavlidis, Jesse Gillis

## Abstract

**Summary:** Evaluating gene networks with respect to known biology is a common task but often a computationally costly one. Many computational experiments are difficult to apply exhaustively in network analysis due to run-times. To permit high-throughput analysis of gene networks, we have implemented a set of very efficient tools to calculate functional properties in networks based on guilt-by-association methods. **EGAD** (**E**xtending ‘**G**uilt-by-**A**ssociation’ by **D**egree) allows gene networks to be evaluated with respect to hundreds or thousands of gene sets. The methods predict novel members of gene groups, assess how well a gene network groups known sets of genes, and determines the degree to which generic predictions drive performance. By allowing fast evaluations, whether of random sets or real functional ones, **EGAD** provides the user with an assessment of performance which can easily be used in controlled evaluations across many parameters.

**Availability and Implementation:** The software package is freely available at https://github.com/sarbal/EGAD and implemented for use in R and Matlab. The package is also freely available under the LGPL license from the Bioconductor web site (http://bioconductor.org).

**Supplementary information:** Supplementary data are available at *Bioinformatics* online and the full manual at http://gillislab.labsites.cshl.edu/software/egad-extending-guilt-by-association-by-degree/.

## 1 Introduction

The analysis of gene networks has emerged as a central interest in bioinformatics with one major focus being to use gene networks to determine commonalities in known gene sets or predict new members (Sharan, et al., 2007). Exploring factors affecting performance is generally so challenging that it is conducted across laboratories, with each group using different algorithms, data, or both (e.g., CAFA (Radivojac, et al., 2013)). Although there are numerous methods to prioritize candidate genes or interpret network data, none are designed to permit controlled computational experiments. Generally, network inference tools benchmark their task on other algorithms or provide basic ‘one-off’ statistical tests (e.g., see (Bellot, et al., 2015; Meyer, et al., 2008)). Instead of exploring the basis of performance in greater detail, a major focus has been in developing more sophisticated algorithms, exacerbating the difficulty of interpretation and controlled comparisons. This focus is surprising in light of the high performance of basic methods in gene network analysis in every controlled evaluation (Consortium, 2014; Eduati, et al., 2015; Gillis and Pavlidis, 2013; Peña-Castillo, et al., 2008). Indeed, simple methods are often generally effective in machine learning (Hand, 2006), performing almost on par to the more “state-of-the-art” approaches. In a series of papers, we have explored factors affecting performance (Gillis and Pavlidis, 2013; Pavlidis and Gillis, 2013; Verleyen, et al., 2015). For example we highlighted the degree to which gene annotation patterns (“multifunctionality”) and network properties influences predictions; e.g., *P53* tends to be a member of many gene sets, which makes it easier to make predictions about its function.

Motivated by these observations, and desiring to make our approaches more widely available, we have developed EGAD (Extending ‘Guilt by Association’ by Degree), a gene network analysis toolset that allows efficient evaluation of thousands of gene sets with respect to the underlying gene network, and also to assess gene properties such as multifunctionality or network topology. EGAD allows the user to determine how well a gene network will group any sets of genes and what genes it would predict as new members of those sets. The methods in EGAD are fully-vectorized and thus orders of magnitude faster than other supervised algorithms. It also assesses the biases and confounds as described in our previous work (Pavlidis and Gillis, 2013). While current offerings include generic graph analysis packages (Csardi and Nepusz, 2006), and packages focused on unsupervised approaches for gene co-expression network analysis (Langfelder and Horvath, 2008), implementations of supervised approaches for functional prediction from gene networks appear to be absent. In addition to the R/Bioconductor package, we provide a Matlab implementation (https://github.com/sarbal/EGAD).

## 2 Implementation and usage

Two of the core features of EGAD are: a function prediction algorithm, allowing network characterization across thousands of functional groups to be accomplished in minutes in cross-validation, and an analytic determination of the optimal prediction across multiple functional sets. We also provide methods to build gene networks from genome-wide biological data sources (e.g., protein-protein interactions).

### Gene networks and gene annotation sets

The network analysis within EGAD require two types of data: a network represented as a matrix, and an annotation set represented as a binary vector. We define networks as undirected graphs, where nodes represent genes and edges the relationship between a pair of genes. For binary networks, edges are either 1s or 0s, indicating whether or not a relationship exists between the adjacent nodes *(build_binary_network)*. These sparse binary networks can be extended (*extend_network)* through indirect connections (Gillis and Pavlidis, 2011), adding weights to the 0s. Co-expression networks (also weighted) can be constructed from the correlation coefficients of gene expression profiles across multiple samples (*build_coexp_network*).

Annotation sets are defined as gene sets, typically corresponding to a given function (e.g., GO: “carbohydrate catabolic process”), a known pathway (e.g., KEGG: “mTOR signaling”), or a disease (e.g., candidate genes for autism). The labels define functional groups represented in an annotation matrix, where rows correspond to genes and columns to the respective groups. Associations between genes and groups are indicated by an entry 1 in the matrix, otherwise are set to 0 (*make_annotations*).

### Neighbor-voting for network analysis and gene function prediction

The neighbor-voting algorithm provided is a fully vectorized method for evaluating functional properties of an interaction network (*neighbor_voting*). It is based on the “guilt-by-association” principle: genes with shared functions are preferentially connected. The algorithm takes a gene network and a set of annotations (gene label vectors) as input. Performed as an *n*-fold cross-validation task, we hide a subset of gene labels and then ask if the remaining genes in the functional group can predict the hidden genes’ identities, in this case having the same annotation (i.e., label) as the selected subset using information inferred from the network. The output is a performance metric for each of the annotation sets tested (often GO functions), and is the averaged AUROC (area under the ROC curve) for each group across the *n*-folds. Analytically calculated for the sake of computation time, this metric broadly reflects how well the genes in the network align with the given annotation set. High AUROCs indicate genes within a gene set preferentially have one another as neighbors.

An additional informative assessment of the network is based on node degree (the number of connections a gene has within a network). Ranking genes according to their node degree, we use that single vector as a prediction for each annotation set. The resulting AUROCs is a measure of node degree bias(Gillis and Pavlidis, 2011), as performance demonstrates the predictability of the gene in the group tested solely due to its prominence in the network rather than its preferential connectivity to other genes within the annotation set.

### Multifunctionality assessment

Another measure to test for biases which can explain performance trends is through a multifunctionality evaluation, based on frequency of annotations across the multitude of sets assessed. Here, a single gene list is constructed which maximizes the performance metric, i.e., gives the highest average AUROC across all functions (*calculate_multifunc, auc_multifunc*). The more highly annotated – multifunctional - a gene is the higher the chances it will be predicted as a good candidate for having any annotation, rendering its prediction correct but uninformative (Gillis and Pavlidis, 2011). A comparison of these multifunctionality AUROCs to the neighbor-voting performance AUROCs gives an indication of the degree to which generic predictions dominate results. Correlation between artifacts and multi-functionality will also induce good apparent performances, often arising due to selection biases (see use case).

### Run time comparisons

To compare the performance of our implementation to other methods, we ran the neighbor-voting algorithm (in R) on a dense co-expression network (17,293 genes) for each GO group of size 20 (104 terms, GO Dec 2014). This took approximately 116 seconds on our server (8 CPU cores @ 2.40GHz). Our implementation in Matlab took ~0.78 seconds. In comparison, the Matlab implementation of GeneMANIA (Warde-Farley, et al., 2010) took 4.2 hours, with highly similar AUROC performance (**Table 1**). To optimize within R, we can use MRAN (https://mran.revolutionanalytics.com/) to obtain a speed-up to approximately 30 seconds.

**Table 1.**
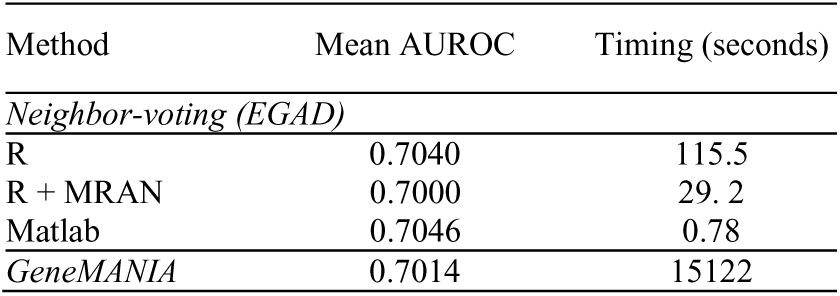
Benchmark results

| Method | Mean AUROC | Timing (seconds) |
| --- | --- | --- |
| *Neighbor-voting (EGAD)* |  |  |
| R | 0.7040 | 115.5 |
| R + MRAN | 0.7000 | 29.2 |
| Matlab | 0.7046 | 0.78 |
| *GeneMANIA* | 0.7014 | 15122 |

### Use case: assessing selection biases in the use of model organisms

It is common for researchers to use gene networks from model organisms in the hope of identifying patterns relevant to humans (Ideker and Sharan, 2008), and thus might consider only genes which have human orthologs. EGAD allows us to explore the biases that might be introduced by this filtering. If we exclude all annotations assigned to genes without orthologs, we are looking at the degree to which GO groups would cluster in gene network data if all functions could only have been originally discovered in mouse, for example. This is a selection bias; the subset of genes without orthologs are now poor candidates for all functional groups. To measure this effect using EGAD, we first constructed a human gene network using BIOGRID (3.4.126), and the corresponding GO annotations matrix. We then created filtered versions of the human annotation matrix limited to the orthologs of each of five model organisms. Then, for each filtered annotation matrix, we calculate a ranked optimal list, which we use to calculate the multifunctionality AUROCs. Additionally, we run the neighbor-voting algorithm (always using the same human gene network) with the filtered annotation matrix, and use the node degree AUROCs from the output.

As shown in Figure 1A and C, limiting analysis to genes having orthologs causes a positive shift in node degree AUROC shift, indicating the filtering selects more well studied and thus more-connected genes. This annotation bias is directly revealed by the multifunctionality assessment (Figure 1B and D): multifunctionality performance (bias) rises as we limit gene function discovery to properties discoverable only in more and more distantly related organisms We conclude that studies of human gene orthologs are more prone to biases in annotations with respect to commonly used network data. In real use-cases, functions arise from many different model organisms, and thus trivially filtering or controlling for these effects will not be possible. Full details and code for this and other use cases are in the user manual.

**Fig. 1.**
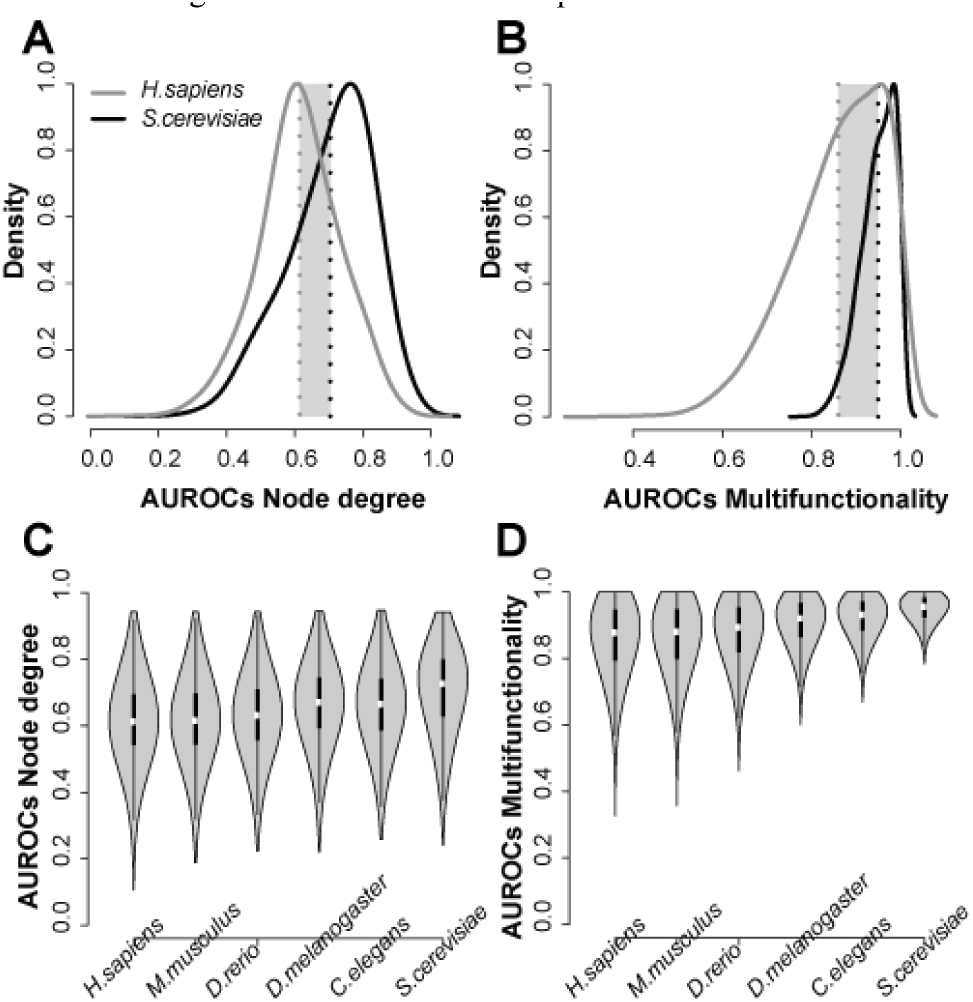
The effect of only considering orthologs as candidates in predicting gene function. AUROCs are the reported average performances for all GO groups evaluated. (A) Node degree performance shift between human (*H. sapiens*, grey) and human genes having yeast orthologs (*S. cerevisiae*, black) (B) and change in multi-functionality performance. (C) The trends across species are consistent for both node degree (C) and multifunctionality (D).

## 3 Discussion

The methods in EGAD are very simple, albeit carefully optimized for efficiency. This is in contrast to the trends in the field to introduce ever more “sophisticated” gene network algorithms, in hopes of exploiting purportedly more complex patterns. Instead, we have shown that most performance seems to be accountable by neighbor-voting, after some simple encoding of global state (i.e., path length to all pairs). This is beneficial because the EGAD methods permit simpler biological interpretations of the networks without sacrificing performance or robustness. We hope that EGAD will encourage researchers performing network-based analysis to thoroughly examine the sources of performance in their own studies and lead to more useful function predictions in the future.

